# Distinct classes of 21 and 24-nt phasiRNAs suggest diverse mechanisms of biogenesis and function in rice anther development

**DOI:** 10.1101/2025.03.03.641266

**Authors:** Rachel Jouni, Caroline Henry, Sébastien Bélanger, Patricia Baldrich, Blake C. Meyers

## Abstract

PhasiRNAs (phased small interfering RNAs) are a major class of plant small RNAs known to be key regulators in male reproductive development of maize (*Zea mays*) and rice (*Oryza sativa*), among other plants. Earlier research focused primarily on premeiotic 21-nt phasiRNAs and meiotic 24-nt phasiRNAs, while new studies uncovered a premeiotic class of 24-nt phasiRNAs. The biogenesis and function of these phasiRNAs remain unclear. We conducted an integrative analysis combining small RNA (sRNA) sequencing and transcriptomic profiling of sRNA-associated genes across ten developmental stages of anther in Kitaake rice to map associations between sRNA-related genes and phasiRNA classes. We identified previously undescribed classes of postmeiotic 21-nucleotide (nt) and 24-nt phasiRNA-producing loci and characterized their unique accumulation patterns. Additionally, our findings reveal distinct nucleotide composition and register accumulation among the phasiRNA classes, suggesting the presence of diverse mechanisms of biogenesis and function. Our results provide new insights into the regulatory complexity of phasiRNAs, establishing a foundation for further functional studies and advancing our understanding of their roles in anther development and their underlying mechanisms.

**Plain Language Summary:** Small RNAs play important roles in controlling how plants develop, but their function in plant reproduction is still not well understood. In rice, small RNAs called phasiRNAs help regulate how pollen develops, but past research has mostly focused on just two types: one found before meiosis and one found during meiosis. In this study, we analyzed small RNA activity across ten stages of rice anther development and found new phasiRNA-producing genes that are active after meiosis. We also discovered that different types of phasiRNAs have distinct patterns of accumulation and composition, suggesting that they have specialized roles in reproductive development. These findings reveal new layers of complexity in how small RNAs function and set the stage for future research to understand their exact roles in pollen formation. This work may help improve our understanding of plant fertility and contribute to advancements in crop breeding.

**Core ideas:**

- Stage-specific accumulation patterns suggest more specialized functions of phasiRNAs.
- Meiotic 24-nt phasiRNAs are compositionally distinct from other 24-nt phasiRNAs.
- Newly described phasiRNA clusters may differ in mechanism of biogenesis and function.

## 1 INTRODUCTION

In order to meet the demands of a growing global population, crop output must increase in a time of growing world hunger, resource limits, and a changing climate. Rice (*Oryza sativa*) is a staple food crop estimated to make up 18.9% of human global caloric intake (“SOFI 2019 - The State of Food Security and Nutrition in the World,” n.d.). Though it is a crop, it is also a model organism for grasses—including wheat and maize–which together with rice account for 42.6% of human global caloric intake (Savary et al. 2019). Breeding crops to improve agronomic traits requires a comprehensive understanding of their genetic makeup and underlying biological mechanisms. The primary focus has been on protein-coding genes, identifying allele variants associated with the most favorable phenotypes, though a common drawback of this approach is unintended phenotypes. Understanding and incorporating gene regulation into breeding efforts may be key to improving crops with fewer negative effects, such as increasing grain yield and engineered pathogen resistance without vegetative growth penalties (Xu et al. 2017; Xiaoguang Song et al. 2022).

One of the major and widespread regulatory mechanisms in plants is the regulation of gene expression by small RNAs. Small RNAs (sRNAs) are 20 to 24 nucleotide (nt) molecules known to regulate genes in a transcriptional (TGS) or posttranscriptional (PTGS) gene-silencing manner. In plants, sRNAs are involved in various developmental processes such as plant development, responses to biotic and abiotic stress, and silencing transposable elements (Axtell 2013a; Bologna and Voinnet 2014). They are typically classified in two main categories: microRNAs (miRNAs) and small interfering RNA (siRNAs). miRNAs originate from single-stranded transcripts produced by RNA polymerase II (Pol II) that form hairpin-like structures that are cleaved by a ribonuclease III enzyme called Dicer-like 1 to produce miRNAs of typically ∼21-nt (Borges and Martienssen 2015). miRNAs principally work in PTGS by guiding the cleavage of target transcripts by Argonaute (AGO) proteins (Borges and Martienssen 2015). In contrast, siRNAs are subdivided into two classes, heterochromatic siRNAs (hc-siRNAs) and secondary siRNAs (Axtell 2013a). hc-siRNAs are derived from precursors transcribed by Pol IV in heterochromatic genomic regions, converted into double-stranded RNA (dsRNA) by RNA-directed RNA Polymerase II (RDR2), and processed into 24-nt sRNAs by DCL3 (Axtell 2013a). Secondary siRNAs include phased secondary siRNAs (phasiRNAs), which originate from Pol II protein-coding transcripts or long noncoding RNA (Fei, Xia, and Meyers 2013; Komiya 2017; Y. Liu et al. 2020). PhasiRNA biogenesis initiates through miRNA-directed, AGO-catalyzed cleavage of single-stranded precursors, which are then converted into dsRNA by RDR6 and processed into 21-nt or 24-nt sRNA duplexes by a DCL protein (Y. Liu et al. 2020).

While 21-nt phasiRNAs are involved in both vegetative and reproductive developments, so far, 24-nt phasiRNAs have only been observed in reproductive development (Teng et al. 2020). PhasiRNAs expressed during reproductive development are named reproductive phasiRNAs and are typically classified in groups of 21-nt premeiotic and 24-nt meiotic phasiRNAs. The 21-nt phasiRNAs were found in seed plants, while 24-nt phasiRNAs were reported only in flowering plants (Pokhrel et al. 2021; Xia et al. 2019). In rice, the biogenesis of 21-nt reproductive phasiRNAs initiates by miR2118-directed AGO1d-catalyzed cleavage of phasiRNA precursors (*PHAS* precursors) transcripts that are converted into dsRNAs by RDR6, processed into 21-nt sRNA duplexes by DCL4, and loaded into AGO1d or AGO5c effectors to trigger the cleavage of specific genes in a posttranscriptional gene silencing manner (Zhai et al. 2015; Shi et al. 2022; Zhan and Meyers 2023; Song, Wang, et al. 2012; Song, Li, et al. 2012; Komiya et al. 2014). In grasses, AGO1d, miR2275, RDR6, and DCL5 are involved in the biogenesis of meiotic 24-nt phasiRNAs, but their AGO effectors and downstream regulatory mechanisms remain largely unknown (Teng et al. 2020; Shi et al. 2022; Song, Wang, et al. 2012; Komiya et al. 2014). Recently, a group of premeiotic 24-nt phasiRNAs was described in barley (*Hordeum vulgare*) and bread wheat (*Triticum aestivum*), and subsequently identified in many other grasses, including rice (Zhan et al. 2024; Bélanger et al. 2020; 2025). Premeiotic 24-nt phasiRNAs differ from the meiotic group in that they are not triggered by miR2275 and are presumably loaded onto a distinct AGO effector.

In this study, we performed sRNA and mRNA sequencing on anthers at ten developmental stages of the rice cultivar Kitaake (*Oryza sativa* subsp. *japonica*). We identified a group of 24-nt phasiRNA loci (24-*PHAS* loci) expressing phasiRNAs at the postmeiotic stage and another group at both premeiotic and postmeiotic stages, which has not been previously reported in the Nipponbare rice cultivar (*Oryza sativa* subsp. *japonica*). Additionally, we observed two groups of 21-*PHAS* loci expressing 21-nt phasiRNAs at postmeiotic stages, also previously unobserved. We analyzed the nucleotide composition of 21-nt and 24-nt phasiRNAs and found that meiotic 24-nt-phasiRNA class is unique among the other phasiRNA classes, further suggesting that this class of phasiRNAs has a unique function. Combined with RNA-seq accumulation of sRNA-related proteins, we identify key genes for mechanistic analysis, paving the way for further exploration of stage-specific phasiRNA functions in rice anther development.

## 2 MATERIALS AND METHODS

### 2.1 Plant materials and growth

Rice Kitaake wild-type seeds were grown in a greenhouse under conditions of 28C/25C day/night, 50-80% relative humidity, and 14-hour day length. We collected anthers at 11 different lengths 0.2, 0.4, 0.6, 0.8, 1.0, 1.2, 1.4, 1.6, 1.8, 2.0, and 2.5 mm from the first three panicles of each plant. At least ten anthers were collected at each stage and promptly fixed with aldehyde for histological analysis. Anther at all stages except 0.2 mm were collected in pools of 30 (≤1.0 mm) or 20 (≥1.2 mm) for RNA experiments. Samples were harvested in three biological replicates, immediately frozen in liquid nitrogen, and stored at −80°^C^ before RNA isolation.

### 2.2 Anther staging

We prepared anthers as described previously (Zhan et al. 2024; Bélanger et al. 2022). Briefly, freshly harvested anthers were fixed overnight in a solution containing 2% [v/v] paraformaldehyde, 2% [v/v] glutaraldehyde, and 0.1% [v/v] Tween20 in 0.1 M sodium cacodylate buffer at pH 7.4. For heavy metal staining, samples were washed (2 x 30 min in water) and transferred in 1.5% [v/v] osmium tetroxide (#19112; EMS, Hatfield, PA, United States) buffered in 0.1 M sodium cacodylate buffer for 3 h. We washed (2 × 60 min in water) and transferred samples in an unbuffered 1% [w/v] aqueous uranyl acetate at 4°C, overnight. Samples in 1% [w/v] aqueous uranyl acetate were then moved to 50°C for 2 h and washed (2 × 60 min in water). Prior to infiltration and embedment, samples were dehydrated following a series of 30, 50, 70, 80, 90, 100 and 100% [v/v] cold acetone at 4°C for 30 min each. Dehydrated samples were exchanged twice in 100% propylene oxide (#20401; EMS, Hatfield, PA, United States) for 30 min each. Samples were then resin infiltrated with a graded series of 25, 50, 75 and 100% Quetol - NSA Kit (#14640, EMS, Hatfield, PA, United States) in propylene oxide without DMP-30 (#14640; EMS, Hatfield, PA, United States) at room temperature for 24 h each step on a rocking platform to enhance resin infiltration. Subsequently, three overnight 100% resin exchanges were made with DMP-30. Finally, samples were embedded in freshly made 100% Quetol using flat embedding molds (#70900; EMS, Hatfield, PA, United States) and polymerized in an oven for 48 h at 60°C. Embedded tissues were then sectioned at 500 nm using the Leica Ultracut UCT (Leica Microsystems Inc., Wetzlar, Germany). Sections were stained using the Epoxy Tissue Stain solution (no. 14950, Electron Microscopy Sciences) and mounted on a slide for image capture using a Leica DM 750 microscope with a Leica ICC50 HD camera along with Leica Acquire v2.0 software (Leica Microsystems Inc., Wetzlar, Germany). The obtained images were subsequently analyzed using ImageJ (Schindelin et al. 2012).

### 2.3 RNA isolation, library construction, and sequencing

Total RNA was extracted with TRI Reagent (#AM9738, Life Technologies, Carlsbad, CA, United States) following the manufacturer’s instructions. sRNA libraries face bias due to capture by library adapters, so we generated libraries using the RealSeq Ac V2 kit for Illumina sequencing (#500-00048, RealSeq Biosciences Inc., Santa Cruz, CA, United States), which reduces bias via degenerate bases and circularization (Baldrich et al. 2020). sRNA libraries were generated following the manufacturer’s recommendations (10 ng of total RNA, adapter dilution of 1:20, and 16 PCR cycles). sRNA libraries were size selected with an end product of ∼150 nt using polyacrylamide gels. For RNA-seq libraries, total RNA was treated with DNase I (#M0303, NEB, Ipswich, MA, United States) and cleaned via ethanol precipitation(Green and Sambrook 2020). After enriching for mRNA using the NEBNext Poly(A) mRNA magnetic enrichment method (#E7490L, NEB, Ipswich, MA, United States), libraries were constructed using the NEBNext Ultra II Directional RNA Library Prep Kit for Illumina (#E7760L, NEB, Ipswich, MA, United States) following the manufacturer’s instructions for use (25 ng RNA after poly-A selection, adapter dilution of 1:20, 16 PCR cycles). The sequencing was performed with single read 51 and paired-end 200 cycles for sRNA-seq and RNA-seq (paired-end) libraries, respectively. The sequencing was generated on an Illumina NextSeq 2000 instrument (Illumina, San Diego, CA, USA) at the University of Delaware DNA Sequencing and Genotyping Center (Newark, DE, USA).

### 2.4 sRNA data analysis

We preprocessed sRNA reads using cutadapt v3.0 (cutadapt -a TGGAATTCTCGGGTGCCAAGGAACTCCAGTCAC -g GTTCAGAGTTCTACAGTCCGACGATC -u 1 -m 10 -j 0 --max-n 0 -o sequencing_file.fq) and we quality controlled preprocessed reads using FastQC v0.12.1 (Martin 2011; “Babraham Bioinformatics - FastQC A Quality Control Tool for High Throughput Sequence Data,” n.d.). Preprocessed reads were aligned to the Oryza sativa Kitaake genome v3.1 (Jain et al. 2019) using bowtie v2.5.2 (bowtie2 -f -N 0 -k 1 --no-1mm-upfront --no-unal --no-sq --norc) (Langmead and Salzberg 2012). Subsequently, the processed reads were aligned to annotated features of the genome, including the coding sequence (CDS), mature microRNAs (miRNAs), ribosomal RNAs (rRNAs), transfer RNAs (tRNAs), transposable elements (TEs), and trans-acting small interfering RNAs (tasiRNAs). RNAmmer was used to annotate rRNA (Lagesen et al. 2007), tRNAscan-SE was used to annotate tRNAs (Chan et al. 2021), and RepeatMasker v4.1.5 was used to annotate TEs (Smit et al. 2013-2015). Sequences of *Oryza sativa* tasiRNAs were obtained from a previous study by the Meyers Laboratory (Xia et al. 2017). To annotate miRNAs from the Kitaake genome, we aligned our reads with the same software and parameters described above to *Oryza sativa* miRNAs in miRBase v22 (Kozomara, Birgaoanu, and Griffiths-Jones 2019). ShortStack v3.8.5 was used to annotate phasiRNAs with a cut-off phasing score of 35 (ShortStack -nohp) (Axtell 2013b).

Aligned reads were normalized to reads per million mapped reads (RPM). Using these normalized reads, size distributions were created for all features for each developmental stage. For both miRNA and phasiRNA datasets, the three biological replicates were averaged and used to filter the miRNA dataset while only those with an average RPM equal to or greater than 10 were kept for downstream analyses. The datasets were then log_2_ normalized and heatmaps were generated using R package pheatmap v1.0.12 (scale = “row”, cluster_rows = TRUE, cluster_cols = FALSE, clustering_distance_rows = “correlation”, clustering_method = “complete”) (Kolde 2012). Using the cutree function in R, we cut the pheatmap dendrograms at heights of 1.75 (Fig. 2b), 1.7 (Fig. 3c), and 1.2 (Fig. 3d) to form expression clusters (R v4.2.0).

**Figure 1.**
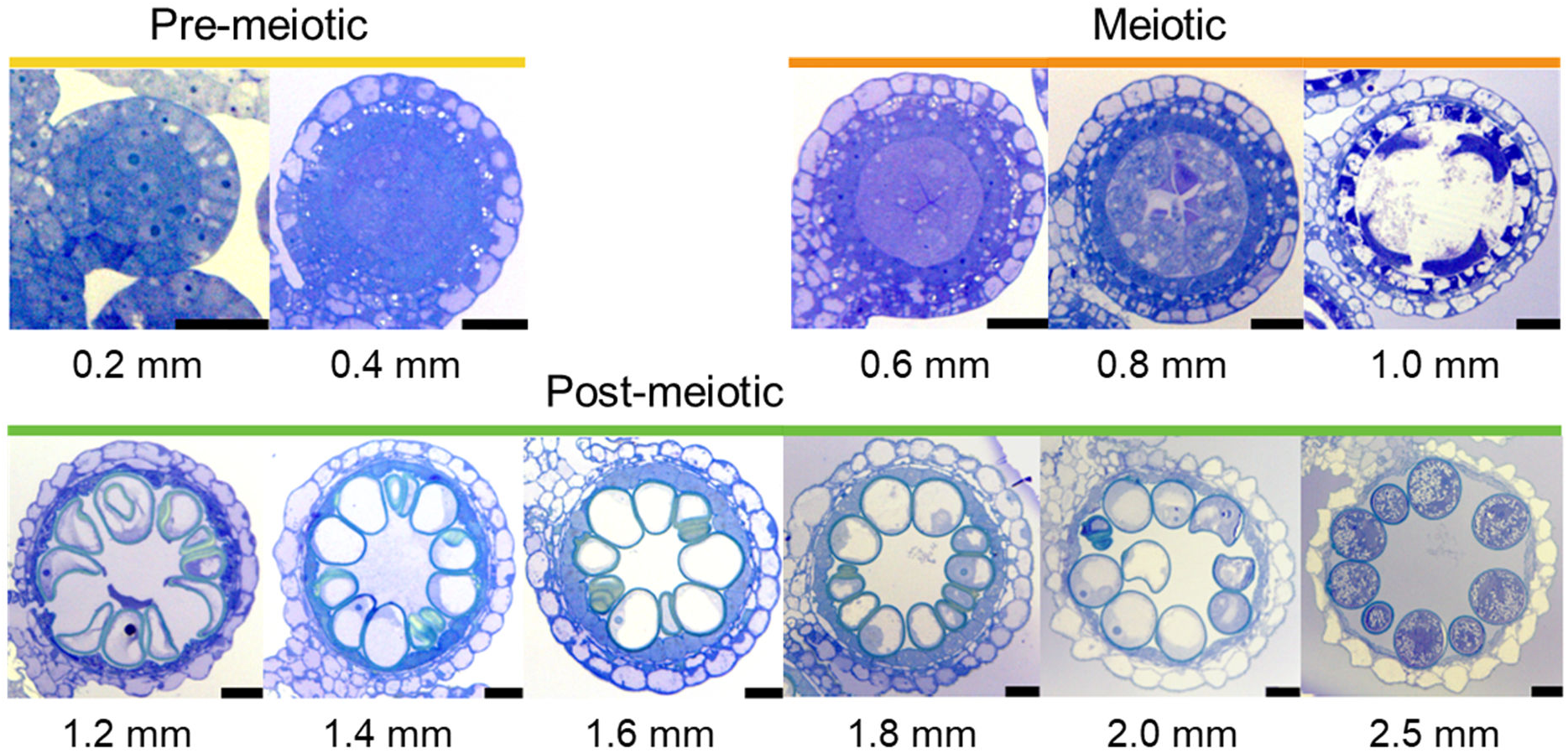
Transverse sections of anthers at 11 developmental stages in Kitaake rice (O. sativa subsp japonica) Anthers sections were taken from the middle of anthers with one lobe imaged. The numbers below each image represent the anther length. The sections with yellow bars represent pre-meiotic stages, the sections with orange bars represent meiotic stages, and the sections with green bars represent post-meiotic stages. Anthers were fixed with a 2% paraformaldehyde:glutaraldehyde solution and embedded using Quetol, sectioned to 500 nm and stained using the Epoxy Tissue Stain solution. Scale bars = 20 µm.

**Figure 2.**
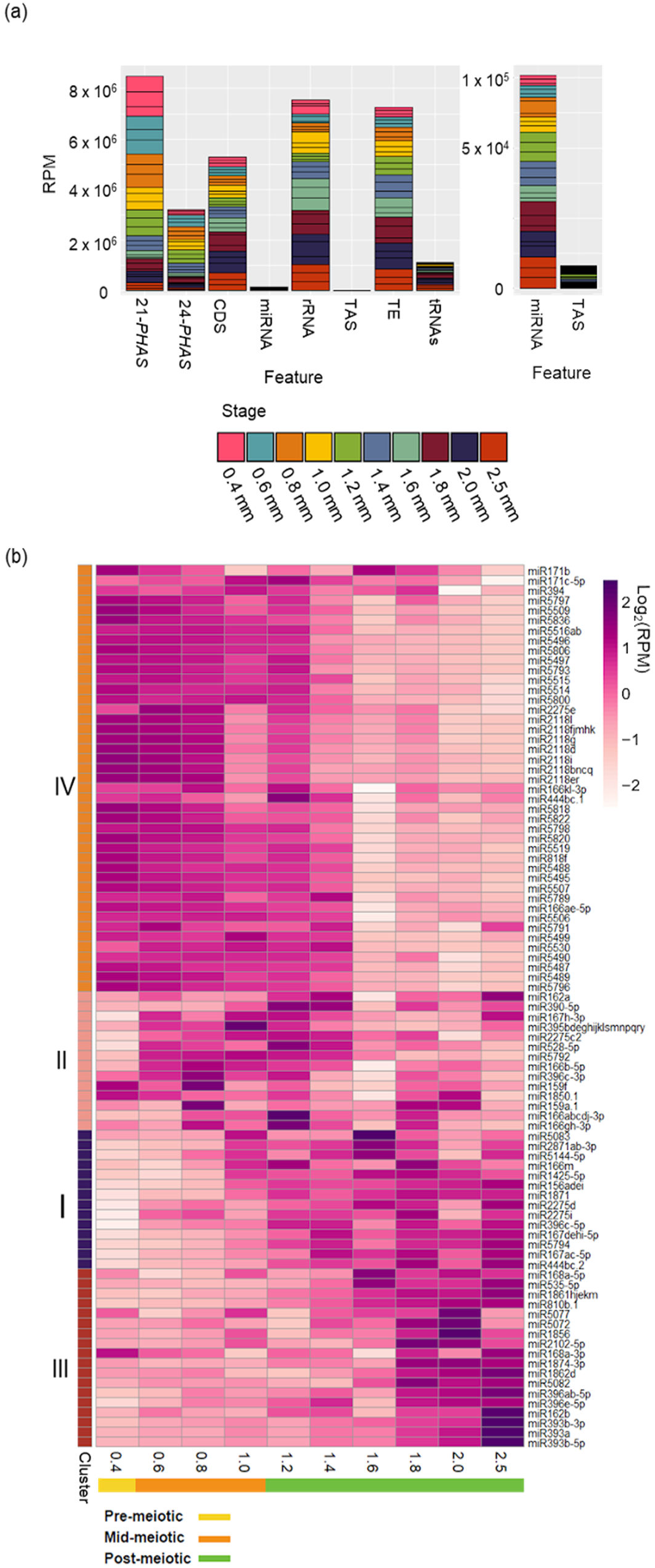
PhasiRNAs are the most abundant sRNAs in rice anther development. (a) Genomic origin of small RNA reads based on genomic features annotated in the Kitaake rice genome v3.1. Mapped sRNA reads that were categorized by genomic features and normalized in reads per million mapped reads (RPM) are represented by stacked bars. The x-axis represents the different features, with the left panel including all features while the right panel represent miRNA and *TAS* loci at a different y-axis scale to enable the visualization. The different colors represent the three biological replicates of each anther stage, from 0.4 mm to 2.5 mm in length. Abbreviations: 21-*PHAS*, 21-nt phasiRNAs; 24-*PHAS*, 24-nt phasiRNAs; CDS, coding sequences; miRNAs, microRNAs; rRNAs, ribosomal RNAs; TAS, trans-acting siRNAs; TEs, transposable elements; tRNAs, transfer RNAs. (b) Heatmap of miRNA expression (log2(RPM)) across anther developmental stages where white represents low accumulation and purple and dark pink represent high accumulation of miRNA reads. Biological replicates were averaged. The numbered bar along the y-axis represents the accumulation clusters, and the bar along the x-axis indicates progression through anther development.

**Figure 3.**
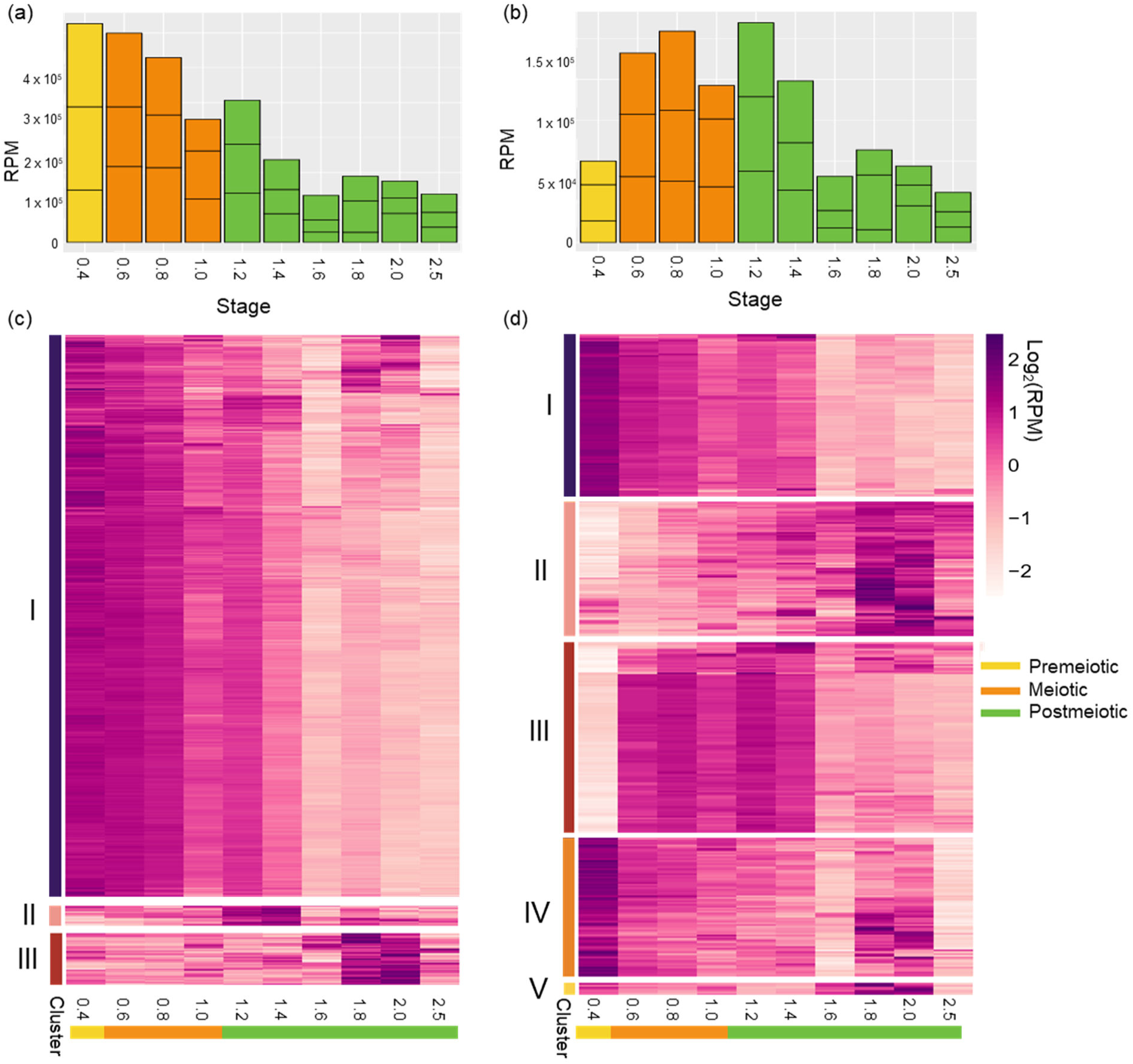
21-nt and 24-nt *PHAS* loci have distinct accumulation patterns. Reads mapping to 21-nt *PHAS* (a) and 24-nt *PHAS* (b) loci identified by ShortStack phasing analysis across anther developmental libraries (supplemental table 2). The x-axis represents the different anther stages, and the y-axis represents the abundance in reads per million (RPM). Data from three independent biological replicates are stacked together in a single bar plot and color-coded by anther stage. (c and d). Heatmaps of phasiRNAs (log2(RPM)) mapping to 21-*PHAS* (c) and 24-*PHAS* (d) loci, scaled by row. Of the three 21-nt *PHAS* clusters, Cluster I contains 1329 loci, Cluster II contains 51 loci, and Cluster III contains 124 loci. Of the five 24-nt *PHAS* clusters, Cluster I contains 97 loci, Cluster II contains 80 loci, Cluster III contains 113 loci, Cluster IV contains 82 loci, and Cluster V contains 7 loci.

### 2.5 Motif enrichment analysis

To identify miRNA target sites in putative *PHAS* precursors, we extended *PHAS* loci coordinates using bedtools slop with a -b value of 500 bp and used it to extract putative precursor sequences using bedtools getfasta (Quinlan and Hall 2010). The extracted precursor sequences were analyzed using MEME Suite v5.5.7 to detect conserved nucleotide motifs with the MEME program (E-value < 0.05) and predict miRNA targeting of identified motifs using the Tomtom program (E-value < 0.001)(Bailey et al. 2015).

### 2.6 phasiRNA analysis

We determined the register by dividing the start position of each read along each locus by the length of the phasiRNAs produced from that locus, either 21 or 24. The remainder dictated the register, and we corrected for the known two nucleotide 3’ overhang (Tamim et al. 2018; Strapps et al. 2010; Elbashir, Lendeckel, and Tuschl 2001). For each locus, we set the highest accumulated register as register 1. Each subsequent register number represents one nucleotide 3’ shift in initial DCL cleavage.

To analyze sequence base composition, we used all the mapped reads from each locus corresponding to an accumulation cluster as described in Figure 3. We filtered for appropriately sized reads (21nt or 24nt) and then used FastQC to analyze the per-position base percentage. The significance of each position was tested via a chi-squared test with a p-value threshold of 0.05. We generated two control groups, one for the 21-nt phasiRNAs and one for the 24-nt phasiRNAs. These control groups were the highest accumulated reads of the unphased sRNA loci (with a phasing score of less than 15) in our ShortStack results.

### 2.7 Annotation of genes involved in sRNA biogenesis and function

Proteome sequence datasets were retrieved from Phytozome (https://phytozome-next.jgi.doe.gov/). OrthoFinder v2.5.4 was used to identify orthologous proteins between Nipponbare and Kitaake rice. The known locus identifiers of sRNA-related genes from Nipponbare were used to select orthogroups for phylogenetic tree inference. Multiple sequence alignment was performed using Muscle v3.8.31, and aligned protein sequences were converted to nucleotide sequences using PAL2NAL v14 (Edgar 2004; Emms and Kelly 2019; Suyama, Torrents, and Bork 2006; Bélanger, Zhan, and Meyers 2023). The maximum-likelihood phylogenetic tree (Supplemental Figure S2) was inferred using IQ-TREE v2.2.0.3 60,61 with default parameters.

### 2.8 RNA-seq library analysis

For RNAseq libraries, raw reads were mapped to the Kitaake reference genome using HISAT2, v2.1.1, and assembled using Stringtie version 2.1.4 (Kim, Langmead, and Salzberg 2015; Shumate et al. 2022). For PCA analysis of our samples, we used DESeq2 v3.2.0 variance stabilized data (Love, Huber, and Anders 2014). For our other analyses, expression levels of genes were normalized on fragments per kilobase million (FPKM) values, and only genes with FPKM > 1 in at least one of the RNA-Seq libraries were kept for downstream analyses. These FPKM values were log_2_-transformed for pheapmap visualization and gene coexpression network analysis using WGCNA v1.72-5 (Kolde 2012; Langfelder and Horvath 2008). We ran WGCNA as previously described by Zhan et al, with certain modifications: (i) the matrix of pairwise Spearman correlation coefficients (SCCs) between genes was transformed into a connection strength matrix by raising the correlation matrix to the power of 22, and (ii) modules with fewer than 100 genes were merged with their closest neighboring modules after hierarchical clustering tree was cut using the Dynamic Tree Cut algorithm to generate the final modules (Langfelder and Horvath 2008; Zhan et al. 2023; Langfelder, Zhang, and Horvath 2008).

## 3 RESULTS AND DISCUSSION

### 3.1 The developmental rate of Kitaake anthers is comparable to that of Nipponbare

Anther development is characterized by a series of sequential cellular differentiation steps yielding specific cell types, and this developmental progression is associated with major shifts in gene expression (Nelms and Walbot 2019; Zhang et al. 2014; Yuan et al. 2018; Deveshwar et al. 2011). The length of anthers is a reproducible phenotypic marker for developmental staging in a given genotype (Marchant and Walbot 2022). Therefore, we performed a cytological analysis of anthers ranging from 0.2 mm to 2.5 mm across 11-time points in the photoperiod-insensitive early-maturity cultivar Kitaake (*Oryza sativa* subsp. *japonica*) to identify key developmental stages (S. L. Kim et al. 2013). We classified anther development into three stages: premeiotic, meiotic, and postmeiotic (Figure 1). At the premeiotic stage, we observed that cell fate specification occurred in anthers measuring < 0.4 mm as a differentiated tapetum was distinguishable at this stage (Figure 1). Meiosis I initiated at 0.6 mm in the meiotic mother cells and meiosis II completed by the 1.2 mm stage where we observed the vacuolated microspores aligned beside tapetal cells, indicating an early postmeiosis stage (Figure 1). The vacuolated microspores develop into mature pollen grains from 1.2 to 2.5 mm when anthesis occurs. Overall, we observed that developmental stages of Kitaake and Nipponbare anthers were similar (Fujita et al. 2010). Based on this staging, we decided to sample anthers at ten developmental stages, from 0.4 mm to 2.5 mm, to provide a comprehensive RNA profiling of Kitaake anther development. The sampling of anthers at six post-meiotic stages, will provide new insights into the pollen maturation expression landscape that had not been covered in previous work in Nipponbare (Fujita et al. 2010).

### 3.2 phasiRNAs dominate the anther developmental sRNA landscape

Previous studies have demonstrated the important roles played by sRNAs to support male fertility in rice and maize anthers (Teng et al. 2020; Shi et al. 2022). To determine the expression pattern of sRNAs in anther development, we sequenced sRNA libraries from 10 anther stages ranging from 0.4 to 2.5 mm in length, with three biological replicates for a total of 30 libraries. We observed that the distribution of read lengths was consistent within biological replicates and differed across the anther stages (Supplementary Figure S1). To determine the genomic origin of sRNA reads, we classified each read by its corresponding locus type (Figure 2a). We observed that the most abundant sRNAs originate from 21-nt phasiRNA loci (21-*PHAS* loci) followed by transposable elements (TEs), ribosomal RNAs (rRNAs), coding sequences (CDS), 24-nt phasiRNAs loci (24-*PHAS* loci), transfer RNAs (tRNAs), and miRNAs. Notably, reads corresponding to miRNAs, TEs, rRNAs, tRNAs, and CDS showed greater accumulation in late postmeiotic stages (1.8 to 2.5 mm), while those overlapping to 21-nt and 24-nt phasiRNA loci (*PHAS* loci) mostly accumulated at earlier stages (Figure 2a). When focusing on reads from *PHAS* loci, the overall abundance of 21-nt phasiRNAs was higher than 24-nt phasiRNAs in anthers at the same stage (Figure 2a). Additionally, premeiotic anthers (0.4 mm) exhibited a high abundance of 21-nt phasiRNAs, whereas 24-nt phasiRNA accumulation was low until the first meiotic stage (0.6 mm) (Figure 2a). These results confirm that phasiRNAs dominate the sRNA landscape in Kitaake anther development.

### 3.3 miRNAs show differing accumulation patterns across anther development

Most of the miRNA annotation work in rice was done in the Nipponbare variety (Jeong et al. 2011; Sunkar et al. 2005; B. Liu et al. 2005; Baldrich, Hsing, and San Segundo 2016). Using annotated rice miRNAs from miRBase v22 (Kozomara, Birgaoanu, and Griffiths-Jones 2019), we identified all of the 582 unique *Oryza sativa* miRNA sequences in our Kitaake anther samples. These miRNAs belonged to 342 different families. We then analyzed the levels of accumulation of each miRNA with an average RPM of 10 (89 miRNAs) or above across all stages (Supplementary Data Set S1). miRNA clustering analysis revealed four expression patterns (Figure 2b). miRNAs in clusters I and III accumulated mostly in late postmeiotic stages (1.8 mm to 2.5 mm), with Cluster III having a higher abundance. Cluster II represents miRNAs expressing at meiotic and early postmeiotic stages (0.6 mm to 1.4 mm), and Cluster IV represents the largest number of miRNA loci, with expression peaking at premeiotic stages and gradually declining through meiosis (0.4 mm to 1.2 mm). The stage of 1.6 mm anthers marks a notable shift in miRNA accumulation, as the miRNAs expressed in premeiotic and meiotic stages (clusters II and IV) show a drop in accumulation at 1.6 mm, while the miRNAs that accumulate in pollen (cluster III) rise in accumulation after the 1.6 mm stage. Our results demonstrate a major shift in the miRNA landscape in postmeiotic stages, separating earlier postmeiotic stages (1.2 mm to 1.6 mm) from later postmeiotic stages (1.8 mm to 2.5mm), a finding made possible by our comprehensive sampling.

The miR2118 family members accumulated in premeiotic stages and through meiosis, consistent with prior reports (Supplementary Data Set 1) (Xia et al. 2019). Interestingly, while two copies of miR2275 showed the expected meiotic accumulation pattern, miR2275d, and miR2275i showed accumulation in later postmeiotic stages (1.6 mm to 2.5 mm). miR168a-3p and miR168a-5p accumulated in late postmeiotic anther development, peaking at 2.5 mm and 1.6 mm, respectively. miR168a-5p is known to regulate AGO activity, and this accumulation pattern was the opposite of previously published *OsAGO1* family expression patterns (Li et al. 2012; Xia et al. 2023). miR168a-3p also peaked in premeiotic stages, suggesting that it may play a role in different stages of anther development. miR162a, known to play a role in balancing resistance to pathogen *Magnaporthe oryzae* and rice yield, also accumulated later in development, peaking at both 1.4 mm and 2.5 mm, while family member miR162b had a strong peak at 2.5 mm (Li et al. 2020). The miR396 family is known to play a role in regulating rice yield, and we observed several copies accumulating in postmeiotic stages (1.8 mm to 2.5 mm). miR396c-3p, however, peaked in meiotic anthers (0.8 mm) (Chandran et al. 2019). The miR393 family is known to function in pollen development, and our results showed a strong peak at 2.5 mm (Jiang et al. 2022). Our profiling of miRNAs showed stage-specific accumulation along anther development, indicating dynamic regulatory roles for miRNAs, with significant shifts from early to late postmeiotic stages.

### 3.4 phasiRNAs show different accumulation patterns corresponding to the main stages of anther development

We hypothesized that the shift in sRNA length that occurred from early stages to late stages corresponds to a shift in phasiRNA accumulation, as this would be consistent with earlier reports (Zhai et al. 2015; Zhan et al. 2024; Fei et al. 2016). We identified 1,544 21-*PHAS* and 379 24-*PHAS* loci encoded in the Kitaake genome (Axtell 2013b) (Supplementary Data Set S2). The number of 21-*PHAS* loci was lower than in past studies (1,544 vs 1,843) in Nipponbare, while the number of 24-nt *PHAS* loci (379 vs 50) was higher, likely due to sampling (Zhan et al. 2024; Fei et al. 2016). For the 21-*PHAS* loci, it is possible that several loci may have been missed in earlier stages than those that we sampled. For the 24-*PHAS* loci, our extra loci may be due to more comprehensive sampling at earlier and later stages in addition to the sampling of anther rather than spikelet in previous work, increasing the sensitivity of our annotation effort. Additionally, previous numbers were based on spikelet tissue rather than dissected anther tissue (Fei et al. 2016). Overall, reads mapping to 21-*PHAS* loci were most abundant in premeiotic anthers and decreased through the end of meiosis, while the burst of reads mapping to 24-*PHAS* loci occurs during meiosis, consistent with previous studies (Figure 3a and c) (Zhai et al. 2015; Fei et al. 2016; Zhan et al. 2024). However, unlike previous studies, the reads mapping to 24-nt *PHAS* loci maintained a steady level of high abundance throughout and after meiosis (0.6 mm to 1.4 mm) (Figure 3b and d) (Zhai et al. 2015).

Assessing the accumulation of the 21-nt phasiRNAs, we observed three patterns or clusters. Cluster I is a premeiotic cluster with a peak accumulation in early stages (0.4 to 1.6 mm), which likely encompasses the premeiotic 21-nt phasiRNAs previously described (Figure 3c). Cluster II and III respectively accumulate at early postmeiotic (1.2 mm and 1.4 mm) and late postmeiotic (1.8 mm to 2.0 mm) stages (Figure 3c). These accumulation patterns had been previously reported in other grasses such as oats, barley, wheat, and rye (Bélanger et al. 2025). The stage specificity of these clusters suggests that the postmeiotic 21-nt phasiRNAs may have a distinct biological function than the premeiotic 21-nt phasiRNAs.

The accumulation of sRNAs from 24-nt *PHAS* loci was grouped into five clusters, based on their level of accumulation in each stage (Figure 3d). Cluster I peaked in the premeiotic stage at 0.4 mm and steadily decreased as development progressed, Cluster II accumulated in late post-meiosis (1.8 to 2.5 mm), cluster III accumulated in early meiosis (0.6 mm) and maintained its level until early post-meiosis (1.4 mm), and Cluster IV accumulated both before (0.4 mm) and after meiosis (1.8 to 2.5 mm). This suggests that unlike the phasiRNAs in clusters I, II, and III, phasiRNAs in cluster IV play a role in two different stages of anther development. Cluster V contained fewer than 10 loci and was excluded from further analysis. Previous literature identified and discussed premeiotic and postmeiotic 21-*PHAS* loci and premeiotic, meiotic, and postmeiotic 24-loci in various grass species (Bélanger et al. 2020; Zhan et al. 2024; Bélanger et al. 2025). This study found that the postmeiotic 21-*PHAS* cluster is split into loci that accumulate in earlier postmeiotic stages and loci that accumulate in later postmeiotic stages, an interesting distinction that mirrors the shift in miRNA accumulation from early to later postmeiotic stages. Additionally, we found a cluster of 24-*PHAS* loci that peak in accumulation in both premeiotic and late postmeiotic stages, suggesting a dual function for this cluster. The identification of new *PHAS* clusters, including those with dual accumulation peaks, suggests a more complex regulation and potentially diverse biological functions of phasiRNAs during anther development than originally thought.

### 3.5 PHAS loci have cluster-specific predicted miRNA targets

The biogenesis of phasiRNAs is initiated by the cleavage of *PHAS* precursors by a miRNA. It was previously reported that miR2275 typically triggers the production of meiotic 24-nt phasiRNAs, and miR2118 triggers the production of premeiotic 21-nt phasiRNAs (Zhan and Meyers 2023). To identify the miRNA triggers of the 21- and 24-nt reproductive phasiRNAs identified in our study, we conducted motif enrichment analysis on the different clusters (as shown in Figure 3) of the *PHAS* precursors. Of the 21-nt phasiRNAs, only clusters I and II (premeiotic and early postmeiotic clusters) contained an enriched miR2118-matching motif. Unlike previous studies, we found that cluster III (late postmeiotic cluster) was not enriched in a miR2118-matching motif (Bélanger et al. 2025). Of the five 24-nt *PHAS* precursor clusters, only cluster III, the meiotic cluster, contained an enriched miR2275-matching motif in 87 of the 113 loci. Cluster II has 80 loci, 39 of which were enriched in a miR1439/1436 containing motif and 9 of which were enriched in a miR2120-matching motif. Clusters I and IV were not enriched in a miRNA-matching motif. Most of these miRNAs had very low levels of accumulation, except for miR1436 and miR2120. Both miRNAs showed a pattern of gradual increase in abundance across anther development through pollen maturation, which is interesting since cluster II peaks in expression in later postmeiotic stages (Figure 3d). Cluster V was discarded due to a low number of loci. While there are some new miRNAs predicted to target loci in the different expression clusters, these miRNAs are predicted to target a smaller percentage of the loci within a cluster compared to miR2118 (21-*PHAS*) and miR2275 (24-*PHAS*). Recent studies described an AU-enriched conserved motif in premeiotic 24-*PHAS* precursors that did not match any miRNA in wheat, barely, oat, maize, teosinte, and many other grasses (Bélanger et al. 2020; Zhan et al. 2024; Bélanger et al. 2025). These results suggest that some of the *PHAS* loci do not require a miRNA for the initiation of their biogenesis.

### 3.6 Meiotic 24-nt phasiRNAs predominantly accumulated one single register

*PHAS* precursors undergo a phased cleavage by Dicer-like proteins, with the initial cleavage site determining the register of resulting phasiRNAs, similar in concept to DNA reading frames. For instance, a 24-nt *PHAS* locus would encompass 24 different registers, while a 21-nt *PHAS* locus would encompass 21 registers. For every predicted *PHAS* locus, we quantified the total counts of phasiRNAs (measured in RPM) starting in each of its registers. To visualize the disparities in register accumulation, we depicted the register counts for each *PHAS* locus on a radar plot. Among the 21-*PHAS* loci, there was a noticeable difference between the highest accumulated register (register 1) and the other registers. Additionally, registers 2 and 21 (one shift in initial cleavage from register 1) and registers 10 and 13 (nine shifts in initial cleavage from register 1) also had higher accumulation, similar to previously published patterns of 21-nt phasiRNA accumulation (Tamim et al. 2018) (Figure 4a). Among the 24-nt phasiRNAs, the accumulation of the registers was more equal, with moderately higher accumulations of registers 1 and 13 (Figure 4a). We then grouped the loci by the previously described expression clusters (Figure 3c and 3d) to search for expression cluster-specific patterns of accumulation. The 21-nt phasiRNA expression clusters all had similar patterns. However, we observed that most of the meiotic 24-nt phasiRNA expression clusters had a greater difference in register 1 counts as compared to the other registers. The rest of the 24-nt phasiRNA clusters showed smaller differences (Figure 4b). This suggests that a greater proportion of meiotic 24-nt phasiRNA reads comes from the top accumulated register than for the other expression clusters.

**Figure 4.**
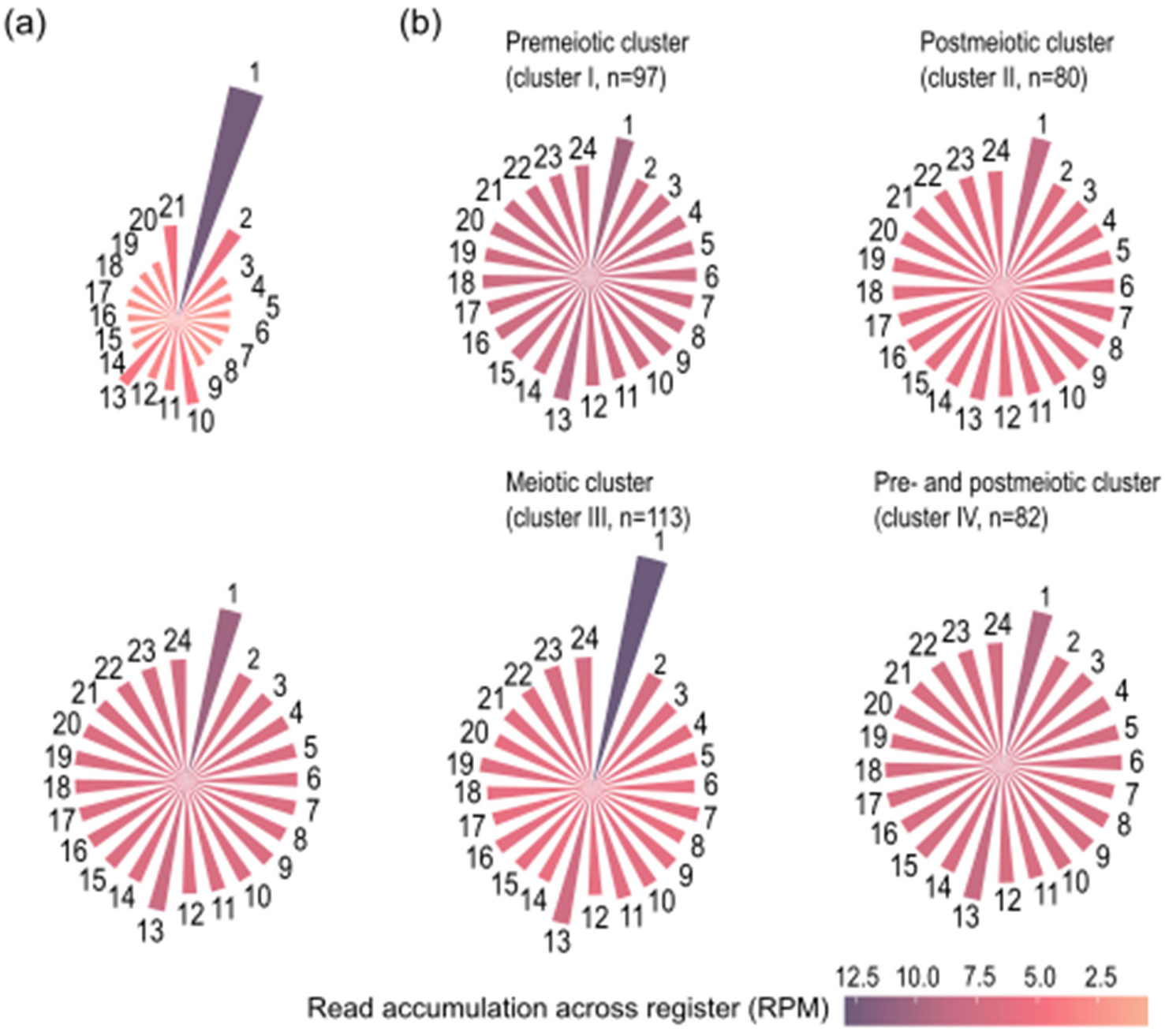
Meiotic 24-nt *PHAS* loci have a pronounced pattern of accumulation by register. (a) The accumulation of reads (RPM) across each register in each 21-nt (top) and 24-nt (bottom) locus. Registers were ordered by nucleotide shift relative to the initial cleavage event, with 1 representing the register with the highest read count (i.e. at the site of initial cleavage), 2 representing one nucleotide shift in the initial cleavage, and so on. (b) The accumulation of reads across each register in the 24-nt *PHAS* loci separated by expression cluster along with the number “n” of loci represented in each cluster.

The register specificity of the meiotic 24-nt phasiRNA cluster compared to the other 24-nt phasiRNA expression clusters has several potential explanations. Our MEME results and previous work show weak evidence for miRNA targeting of non-meiotic 24-*PHAS* loci, suggesting a potential miRNA-independent biogenesis pathway for these loci. This pathway may lead to greater variation in the initial Dicer cleavage site than a miRNA-dependent pathway. Alternatively, it is possible that there are weaker, and possibly multiple, miRNA targets within non-meiotic 24-*PHAS* loci filtered out by our criteria. Another possibility is that multiple Dicer proteins are capable of processing the other expression clusters while only one targets the meiotic cluster. Zhan *et al*. hypothesized that the biogenesis of meiotic 24-nt phasiRNAs are more reliant on DCL5, while premeiotic 24-nt phasiRNAs may be reliant on both DCL5 and DCL3 (Zhan et al. 2024). In this case, if multiple Dicers cleave a locus, then phasiRNAs from different registers are produced. The 21-*PHAS* loci had a second and smaller peak in accumulation one shift away from the highest accumulated register, while the 24-*PHAS* loci did not, suggesting that OsDCL4 shifts its initial cleavage more than DCL5/DCL3, which processes the 24-nt phasiRNA precursors. Both the 21-*PHAS* and 24-*PHAS* loci have peaks that are shifted nine and twelve positions from their highest accumulated register, which is likely the products produced by their self-cleavage (Tamim et al. 2018).

### 3.7 Differential nucleotide preferences indicate cluster-specific 5’ and 3’ nucleotide biases among 24-nt phasiRNAs

Small RNAs are loaded into AGO proteins, a process partially directed by the 5’ nucleotide of the sRNA, and more generally by the sRNA nucleotide composition (Mi et al. 2008). Thus, we investigated the nucleotide composition 21- and 24-nt phasiRNAs originating from *PHAS* loci based on their accumulation pattern described in Figure 3. We observed cluster-specific sequence trends for both the 21- and 24-nt phasiRNAs (Figure 5). For the 21-nt phasiRNAs, clusters II and III (early and late postmeiotic) had much more variation in base pair composition than cluster I, likely due to the fact that these two clusters have much fewer loci. Thus, we focused on cluster I, as it contained the majority of the loci (88%). Our results showed a significant preference for a C in position 1, as described in previous literature. Previous research on combined maize and rice data found that among the 21-nt phasiRNAs, positions 1, 3, 14, 19, and 21 were significant (Patel et al. 2018). Similarly, another study in 7 plant species found C/G enrichment at positions 1, 3, 19, and 21 (C/G), while positions 3, 8, 14, and 20 were (A/U-enriched) (Bélanger et al. 2025). There was a slight preference for G at position 14 instead of U, as previously described, but this was not significant. Our results support the significance of the 5’ nucleotide in premeiotic 21-nt phasiRNAs and suggest the importance of position 14. In our analysis, we used non-phased reproductive phasiRNAs as a control, meaning that our identified significant positions are specific to reproductive phasiRNAs.

**Figure 5.**
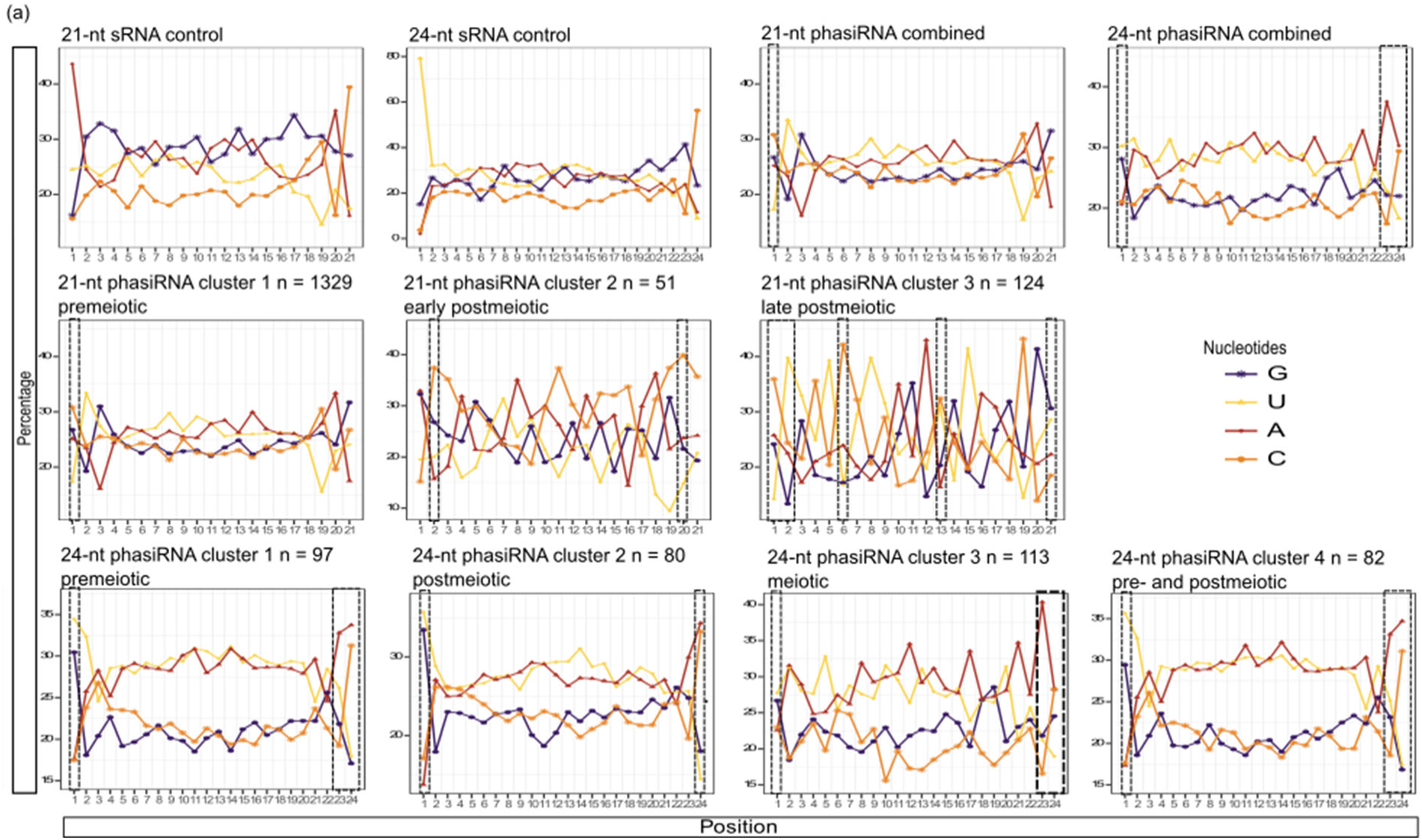
Meiotic 24-nt phasiRNAs in rice do not show a trend in the 5’ nucleotide. Per base sequence composition of phasiRNAs by accumulation cluster. For each locus we took all reads of appropriate length locus and grouped them by accumulation cluster. We then ran them through FastQC. Significance, indicated by the dashed-line boxes, was calculated with a chi-squared test with the 21-nt and 24-nt sRNAs as a control and a p-value of 0.05.

For the 24-nt phasiRNAs, the previous study in maize and rice showed a significant enrichment in A/U nucleotides at positions 1, 10, 20, 21, 22, and 23 (Patel et al. 2018), while the study conducted in seven species found nucleotide biases at positions 1 (A/G), 2 (A), 23 (U), and 24 (U/C) (Bélanger et al. 2025). Our data shows a general bias for A and U across the length of the 24-nt phasiRNAs regardless of cluster. We also observed significantly higher rates of position 1 G/U nucleotides for all clusters except for cluster III, the meiotic cluster, which did not have a dominant 5’ nucleotide. The 24-nt phasiRNA clusters also had higher levels of 3’ G/C reads with the exception of cluster III, in which the difference was smaller. For each cluster except for the postmeiotic cluster II, positions 23 and 24 are significant with preferences for A and C. Interestingly, the cluster with the smallest 3’ nucleotide preference is the meiotic cluster III. Compared to previous studies, our data highlights the importance of the 5’ and 3’ nucleotides, even if base identity is not conserved. It is unexpected that the meiotic 24-nt phasiRNAs do not have an apparent predominant 5’ nucleotide, as this has not been described in previous literature, likely due to a difference in analysis methods, as we used all reads rather than the top accumulated read per locus and we separated the meiotic and post meiotic clusters (Zhan et al. 2024; Bélanger et al. 2025). To best represent phasiRNA biology, we analyzed all reads from a locus as opposed to representing each locus with its highest accumulated read. We also analyzed the composition of meiotic and postmeiotic 24-nt phasiRNAs separately. Since the 5’ nucleotide of an sRNA influences the identity of the AGO that loads it, and previous research has demonstrated AGO loading of meiotic 24-nt phasiRNAs, this result indicates that meiotic 24-nt phasiRNAs are loaded into multiple classes of AGO proteins (Zhan et al. 2024; Mi et al. 2008; Tamotsu et al. 2023). Given both the rise in 24-nt phasiRNA accumulation during meiosis (Figure 3b) and their apparent lack of non-cis targets, meiotic 24-nt phasiRNAs may flood AGOs present during meiosis. Going forward, studies of individual AGO proteins using RNA immunoprecipitation sequencing are necessary to see if the differences observed in 5’ nucleotides translate into differential AGO loading.

### 3.8 Genes encoding small RNA-related proteins display stage-specific expression during anther development

The biogenesis and function of different sRNA classes rely on various small RNA-related proteins such as RNA-dependent RNA polymerases (RDRs), Dicer-like proteins (DCLs), and AGO proteins. In many plants, including rice, specialized copies of these proteins operate in specific sRNA pathways, and identifying the specific proteins associated with a sRNA can elucidate its function (Zhan and Meyers 2023). To explore the expression patterns of genes associated with sRNA biogenesis, we also performed an RNA-seq experiment on rice anthers at the previously mentioned ten developmental stages. Reads were mapped to the Kitaake genome, and we calculated the average fragments per kilobase of transcript per million mapped reads (FPKM) for each gene. Principal Component Analysis (PCA) on DESeq2 variance-stabilized data revealed stage-specific relationships among samples (Figure 6a) (Love, Huber, and Anders 2014). Biological replicates are grouped by developmental stages, with significant PC1 and PC2 distances distinguishing later postmeiotic stages (1.6 mm to 2.5 mm) from earlier stages, suggesting a substantial shift in gene expression during pollen maturation.

**Figure 6.**
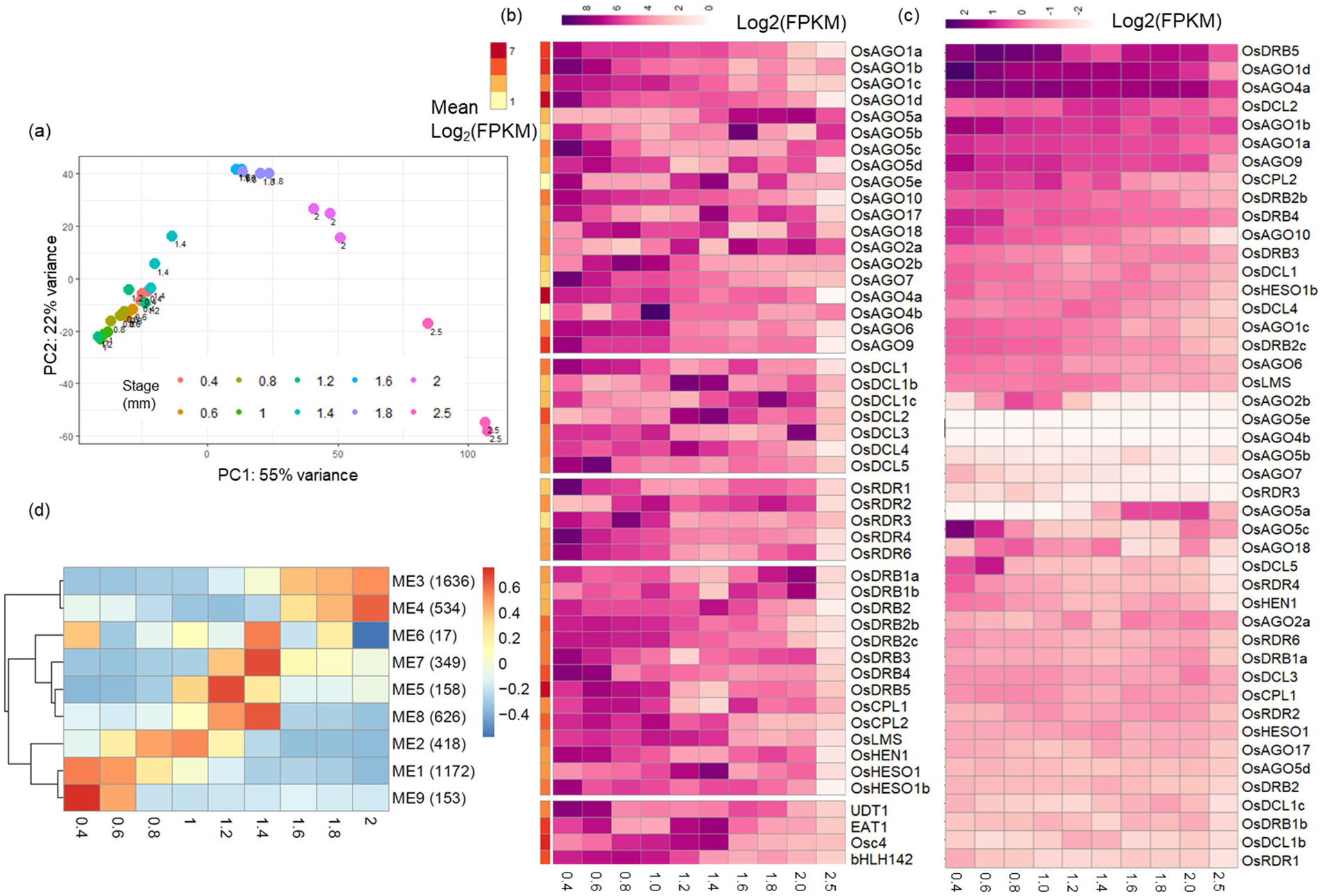
Transcripts of sRNA-related genes show distinct accumulation patterns across gene families and duplications. (a) Principal component analysis of the different anther libraries and their three biological replicates. (b) Heatmap of the transcripts (FPKM) of sRNA-associated genes across anther development without scaling. (c) Heatmap of the transcripts (FPKM) of sRNA-associated genes across anther development scaled by row. (d) Heat map of module eigengenes (MEs) along with number of genes in each module (in parentheses) of the 9 co-expression modules of the annotated genes from Kitaake with the phasiRNA precursors.

To validate our staging, we checked the expression patterns of four genes known to play important roles in anther development. In previous studies, *Undeveloped Tapetum1 (Udt1)*, a major regulator of tapetal development, is shown to have low transcript levels and to express in anthers during earlier meiotic stages (Jung et al. 2005). *ETERNAL TAPETUM1 (EAT1)*, a basic-helix-loop-helix (bHLH) transcription factor, was shown to induce programmed cell death (PCD) in postmeiotic anther tapetum and expresses in both early meiosis and post-meiosis (Ono et al. 2018). *Osc4* was shown to express throughout meiosis and earlier stages of pollen maturation (Fujita et al. 2010). Finally, the bHLH transcription factor *bHLH142* was shown to accumulate during meiosis and decline in postmeiosis and is absent from pollen (Ranjan et al. 2017). Our results reflect these patterns (Figure 6b).

To identify sRNA-associated genes correlated with phasiRNA clusters, we constructed a maximum-likelihood phylogenetic tree (Supplementary Figure S2) to identify homologs of Nipponbare sRNA-associated genes in Kitaake. We then plotted the expression patterns of these genes (Supplementary Data Set S3). Among the OsAGO proteins, OsAGO5c (MEL1), OsAGO1b, and OsAGO1d are known to load phasiRNAs or their miRNA triggers in premeiotic anthers, which is reflected in our results (Figure 6c) (Shi et al. 2022; Komiya et al. 2014; Tamotsu et al. 2023). *OsDRB5* had high expression levels and showed peaks in expression during meiosis and again in later postmeiotic stages (1.6 mm to 2.0 mm). Some proteins, though not as highly accumulated, show more specific patterns of expression. *OsAGO2b* was expressed exclusively during meiosis and shortly after (0.6 mm to 1.2 mm anthers), whereas *OsAGO2a*, a tandem duplication, showed increased expression post-meiosis (1.6 mm to 2.5 mm anthers). Another tandem duplication, *OsAGO5e*, had low expression levels, whereas *OsAGO5a* increased expression in post-meiosis. *OsAGO4a*, involved in RNA-directed DNA methylation, consistently showed high accumulation throughout anther development, and *OsDCL5* expression peaked during early meiosis (Zhan and Meyers 2023). A previous study found an overlap between the accumulation of stage-specific phasiRNAs and sRNA-related protein transcripts (Bélanger et al. 2020). *OsDCL5* and *OsAGO2b* expression overlap with 24-nt phasiRNA Cluster III (meiotic) accumulation, with *OsDCL5* expression aligning with earlier meiotic stages. *OsAGO5a* expression coincided with 24-nt phasiRNA Cluster II and IV (postmeiotic) accumulation. The three copies of *ZmAGO18* are known to be important for the maize 24-nt phasiRNA pathway (Zhan et al. 2024). The single *OsAGO18* copy showed moderate expression overlapping with 24-nt phasiRNA Cluster III accumulation and in 2.0 mm anthers, hinting at its possible role in the rice 24-nt phasiRNA pathway. The stage-specificity of the expression patterns paints these genes as attractive potential targets for studying their corresponding phasiRNA clusters.

When comparing transcript abundance scaled by row, which better demonstrates patterns of gene expression, we observed distinct patterns among *OsDCL* and *OsRDR* family members (Figure 6b). *OsDCL1* showed the highest transcript accumulation in premeiotic anthers, decreasing across development, while *OsDCL5* peaked at meiosis I. *OsDCL1b* and *OsDCL2* peaked post-meiosis, and *OsDCL1c* had moderate premeiotic accumulation with a specific peak post-meiosis. *OsDCL3* accumulated moderately through meiosis, peaking at late-post meiosis. Given these patterns, *OsDCL2* may play a role in the biogenesis of postmeiotic phasiRNAs, while *OsDCL5* and *OsDCL1* overlap with meiotic and premeiotic phasiRNA expression, respectively. Since *PHAS* loci express before the phasiRNAs accumulate, it is possible that there is an OsDCL protein expressed in 0.2 mm anthers that is responsible for the premeiotic phasiRNA clusters. The *OsRDR* transcripts generally decreased in expression across anther development, except for *OsRDR2*, which had moderate expression from 0.8 mm to 2.0 mm anthers, and OsRDR3, which peaked specifically during meiosis II, despite its low accumulation.

To get a sense of how our expression results compared to the overall expression of RNA transcripts across anther development, we conducted WGCNA analysis using our RNA-seq libraries. We identified 9 ME modules with 3 main shifts in expression (Figure 6d) (Supplementary Data Set S3). The first shift in expression is in premeiotic to meiotic stages (from 0.4 mm to 0.6 mm), the second is after meiosis in early postmeiotic stages (from 1.0 mm to 1.2 mm and 1.4 mm) and the last is in later-post meiotic stages (from 1.8 mm to 2.0 mm). The first shift is similar to 24-nt phasiRNA clusters I and IV expression patterns (Figure 3d), as well as the patterns of the premeiotic transcripts such as *OsAGO5c* and *OsAGO1d* (Figure 6b and c). The second shift is similar to 21-nt phasiRNA cluster II expression (Figure 3c) as well as *OsDCL1b*, *OsDCL2*, and *OsAGO18* expression patterns (Figure 6b and c). There is a general decrease in expression at 1.6 mm, which is consistent across miRNA accumulation (Figure 2b), 21- and 24-nt phasiRNA accumulation (Figure 3c and 3d), and the WGCNA expression modules (Figure 6d). The late postmeiotic expression shift is similar to miR393 expression (Figure 2b), as 21-nt phasiRNA Cluster III (Figure 3c), 24-nt phasiRNA clusters II and IV (Figure 3d), and patterns of *OsAGO5a*, *OsDCL1c*, *OsDCL3*, *OsDRB1a*, and *OsDRB1b* (Figure 6b and 6c). Taken together, our analysis provides insight into the dynamics of sRNA-related gene expression throughout anther development in rice. We identified stage-specific expression patterns of key sRNA-related proteins such as OsAGO, OsDCL, and OsRDR family members, highlighting their diversified roles in phasiRNA biogenesis and functions. The correlations between the expression of these genes and the accumulation patterns of 21- and 24-nt phasiRNA clusters suggest complex regulatory mechanisms orchestrating anther developmental sRNA pathways. These findings provide a foundation for future studies on the functional implications of these sRNA pathways and their contributions to plant reproductive development.

## Supporting information

Supplementary Data Set S1

Supplementary Data Set S2

Supplementary Data Set S3

## Abbreviations

AGO: Argonaute
bHLH: basic-helix-loop-helix
CDS: coding sequence
DCL: Dicer-like protein
FPKM: fragment per kilobase million
miRNA: microRNA
PC: principal component
PCA: principal component analysis
PCD: programmed cell death
PTGS: post-transcriptional gene silencing
phasiRNA: phased secondary siRNA
*PHAS* precursors: phasiRNA precursors
RDR: RNA-directed RNA polymerase
RPM: reads per kilobase million mapped
rRNA: ribosomal RNA
siRNA: small interfering RNA
sRNA: small RNA
TAS: trans-acting siRNAs
TE: transposable elements
TGS: transcriptional gene silencing
tRNA: transfer RNAs
WGCNA: weighted gene co-expression network analysis

## ACKNOWLEDGMENTS

We thank Mark Shaw and Brewster Kingham from the University of Delaware DNA Sequencing & Genotyping Center for providing their expertise and assistance in sequencing.

## SUPPLEMENTAL MATERIAL

**Supplementary Figure S1:** size distribution of sRNA libraries

**Supplementary Figure S2:** Phylogenetic tree of Kitaake and Nipponbare sRNA-related genes

**Supplementary Figure S3:** Accumulation patterns of miR2120 and miR1436

**Supplementary Table S1.** Predicted mRNA targets of phasiRNA clusters

**Supplementary Data Set S1:** Counts of sRNA reads mapping to annotated miRNAs

**Supplementary Data Set S2:** Counts of sRNA reads mapping to *PHAS* loci

**Supplementary Data Set S3:** Counts of RNA-seq reads mapping to sRNA-related transcripts and WGCNA module identity

