## Supplemental Material for "Distinct classes of 21 and 24-nt phasiRNAs suggest diverse mechanisms of biogenesis and function in rice anther development"

**Supplementary Figures**

**Page 2: Supplementary Figure S1:** size distribution of sRNA libraries

**Page 3: Supplementary Figure S2:** Phylogenetic tree of Kitaake and Nipponbare sRNA-related genes

**Page 4: Supplementary Figure S3:** Accumulation patterns of miR2120 and miR1436

Page 5: Supplementary Table S1. Predicted mRNA targets of phasiRNA clusters

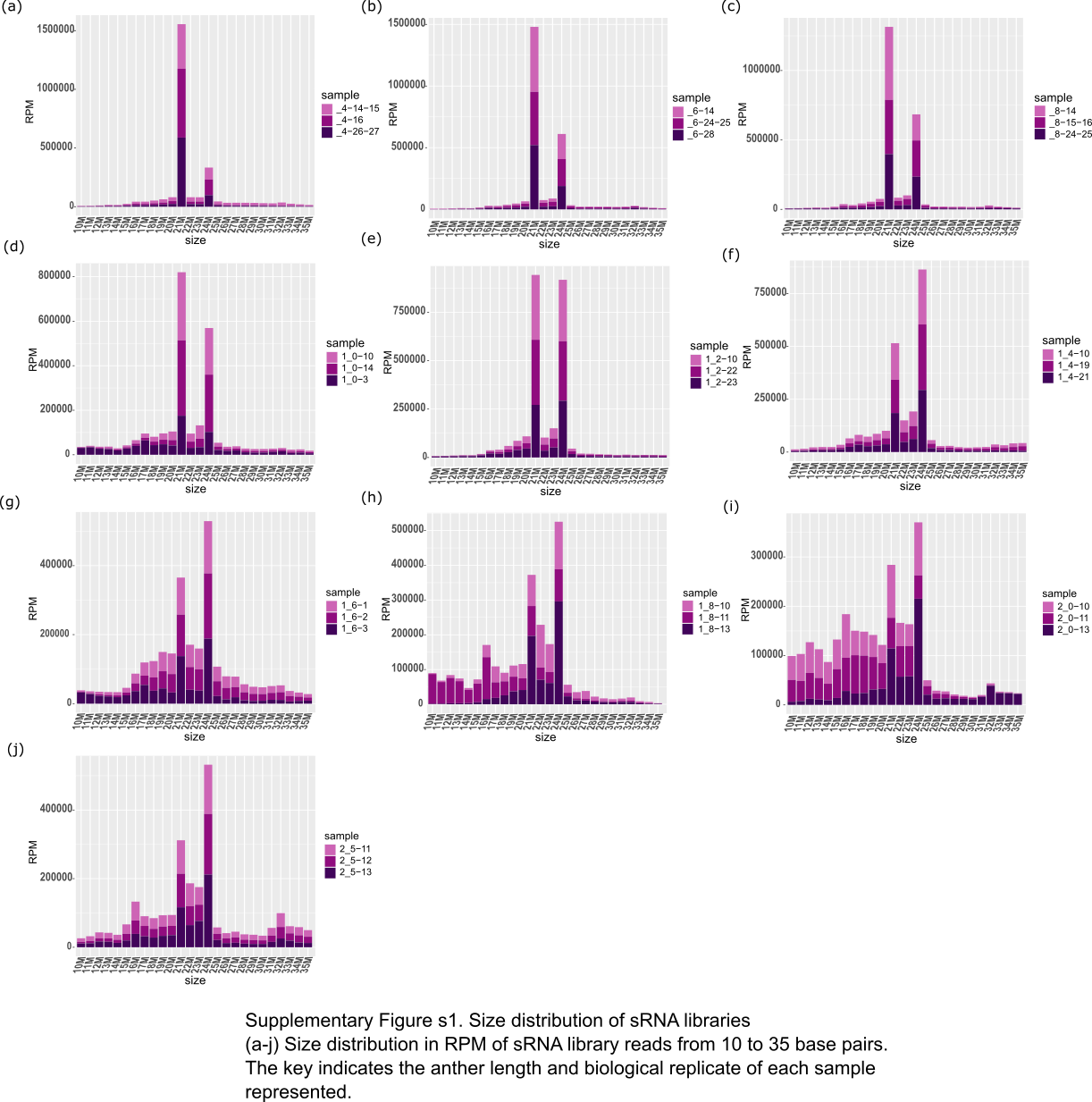
**Supplementary Figure S1. Size distribution of sRNA libraries**

(a-j) Size distribution in RPM of sRNA library reads from 10 to 35 base pairs. The key indicates the anther length and biological replicate of each sample represented.

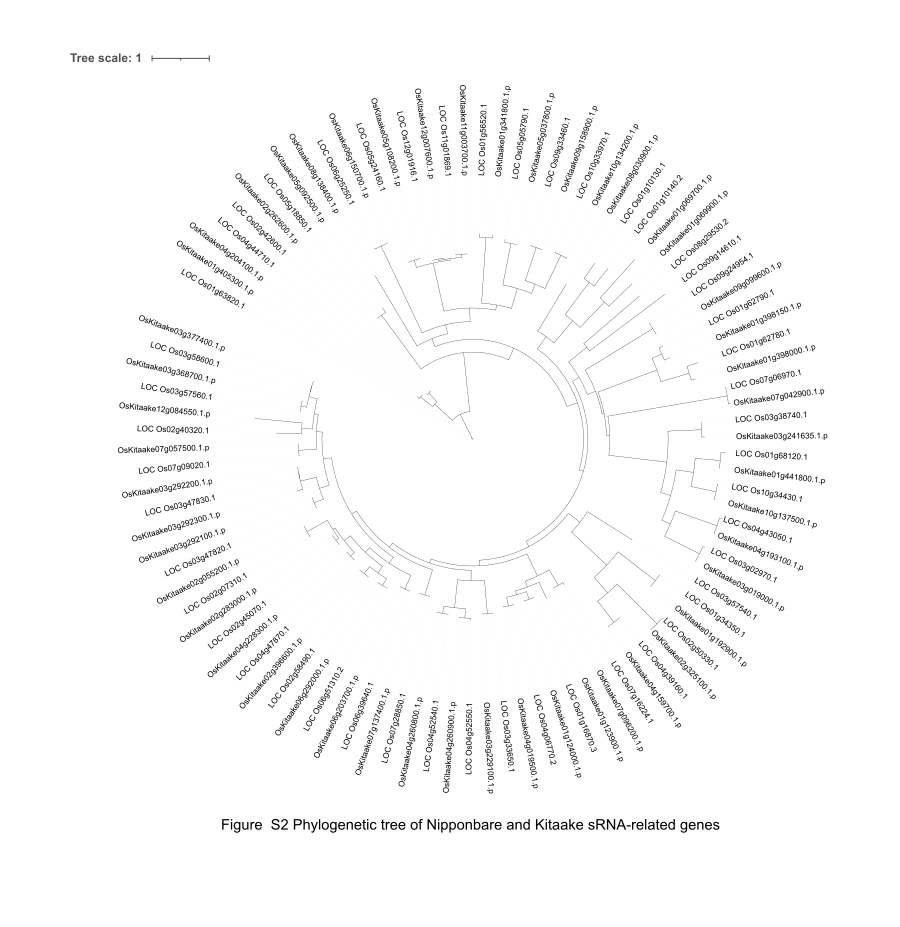

Supplementary Figure S2: Phylogenetic tree of Kitaake and Nipponbare sRNA-related genes

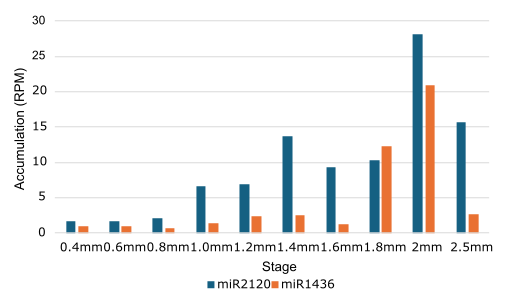

**Supplementary Figure S3:** Accumulation patterns of miR2120 and miR1436

Supplementary Table S1. Predicted mRNA targets of phasiRNA clusters

| Length | Cluster | Predicted miRNA-target | TOMTOM  e-value |
| --- | --- | --- | --- |
| 21 | I (premeiotic) | miR2118 | 7.58 x 10^-11^ |
|  | II (early postmeiotic) | miR2118 | 9.44 x 10^-11^ |
| 24 | II (postmeiotic) | miR1436/1439 | 1.0 x 10^-5^/2.2 x 10-^5^ |
|  | II (postmeiotic) | miR2120 | 1.4 x 10^-7^ |
|  | III (meiotic) | miR2275 | 3.75 x 10^-10^ |
